# REPrise: *de novo* interspersed repeat detection using inexact seeding

**DOI:** 10.1101/2024.01.21.576581

**Authors:** Atsushi Takeda, Daisuke Nonaka, Yuta Imazu, Tsukasa Fukunaga, Michiaki Hamada

## Abstract

**Motivation:** Interspersed repeats occupy a large part of many eukaryotic genomes, and thus their accurate annotation is essential for various genome analyses. Database-free *de novo* repeat detection approaches are powerful for annotating genomes that lack well-curated repeat databases. However, existing tools do not yet have sufficient repeat detection performance.

**Results:** In this study, we developed REPrise, a *de novo* interspersed repeat detection software program based on a seed-and-extension method. Although the algorithm of REPrise is similar to that of RepeatScout, which is currently the de facto standard tool, we incorporated three unique techniques into REPrise: inexact seeding, affine gap scoring and loose masking. Analyses of rice and simulation genome datasets showed that REPrise outperformed RepeatScout in terms of sensitivity, especially when the repeat sequences contained many mutations. Furthermore, when applied to the complete human genome dataset T2T-CHM13, REPrise demonstrated the potential to detect novel repeat sequence families.

**Availability:** The source code of REPrise is freely available at https://github.com/hmdlab/REPrise. Repeat annotations predicted for the T2T genome using REPrise are also available at https://waseda.box.com/v/REPrise-data.

## 1 Introduction

Interspersed repeats, which are mainly amplified by copying of transposable elements (TEs) while undergoing evolutionary mutations, constitute a significant portion of many eukaryotic genomes. For example, they account for 54% of the human genome (Hoyt *et al*., 2022) and 85% of the wheat genome (The International Wheat Genome Sequencing Consortium (IWGSC) *et al*., 2018). These repeat sequences were once considered functionless ‘junk’ regions, but they are now understood to play various roles in cellular processes, including RNA processing and transcriptional regulation (Attig *et al*., 2018; Chishima *et al*., 2018; Zeng *et al*., 2021). Moreover, because these repeat sequences contain phylogenetic signals, they are employed as markers for reconstructing species trees (Dodsworth *et al*., 2015). Consequently, the accurate annotation of interspersed repeats is an essential task of genome analysis.

There are two main computational approaches for identifying interspersed repeats: database-dependent and *de novo* database-free methods. The former annotates the repeats by aligning sequences from repeat databases, such as Repbase (Bao *et al*., 2015) and Dfam (Storer *et al*., 2021), to the genome. The representative tool for this approach is RepeatMasker (Smit, 2015), which is widely used for annotating repeats in newly sequenced genomes (Hoyt *et al*., 2022). However, this approach inherently faces challenges in annotating genomes without well-curated repeat databases and in detecting repeats absent from existing databases. Recent advances in sequencing technology have led to numerous cases that cannot be adequately handled by this database-dependent approach alone. For example, the Earth BioGenome Project (Lewin *et al*., 2022), which aims to sequence all eukaryotic genomes by 2030, continues to sequence the genomes of many non-model organisms that lack sufficient repeat annotation. As another example, the Telomere-to-Telomere (T2T) Consortium has released a complete human genome using long read sequencers and has discovered novel repeat families in newly sequenced regions (Nurk *et al*., 2022; Hoyt *et al*., 2022).

In these cases, the latter *de novo* database-free approach is practical for detecting interspersed repeats. At present, various tools based on this approach have been developed (Bao and Eddy, 2002; Gu *et al*., 2008; Flutre *et al*., 2011; Girgis, 2015; Schaeffer *et al*., 2016; Feng *et al*., 2020), and benchmark studies have indicated that RepeatScout (Price *et al*., 2005) is the best-performing tool (Saha *et al*., 2008; Ou *et al*., 2019; Rodriguez and Makałowski, 2022). RepeatScout employs a seed-and-extension method, originally proposed as a fast alignment heuristic in BLAST (Altschul *et al*., 1990). RepeatScout first identifies frequently occurring seed regions within the genome and then performs extension alignments from both ends of these seed regions. Once the extension alignment is completed, the corresponding seed regions are masked, and the extension alignment is performed again from different seed regions. This cycle of alignment and masking continues until alignments from all seed regions have been performed. Due to its high repeat detection performance, RepeatScout has been incorporated into comprehensive repeat annotation pipelines such as EDTA (Ou *et al*., 2019) and RepeatModeler2 (Flynn *et al*., 2020). However, even RepeatScout has yet to achieve perfect repeat detection, indicating that there is room for further improvement in the algorithm. In addition, due to implementation issues, RepeatScout cannot be applied directly to long genome sequences such as the human genome.

In this study, we developed REPrise (REPeat Recognition using Inexact Seed-and-Extension), a tool that identifies interspersed repeats *de novo* with higher sensitivity than RepeatScout. The algorithm of REPrise is organized into three main steps similar to those in RepeatScout: the seed detection, the extension alignment, and the masking. REPrise improved each of these steps compared to RepeatScout as follows.

1. Unlike RepeatScout, which used exact *k*-mer matches as seed sequences, REPrise employed inexact seeds that allow for *d* mismatches. Because some highly sensitive sequence alignment tools utilized seeding that are not exact matches (Ma *et al*., 2002), we expected that the inexact seeding improves the detection sensitivity of interspersed repeats. Note that phRAIDER (Schaeffer *et al*., 2016) used seeding that are not exact matches for interspersed repeat detection, but phRAIDER is limited in the detectable repeat types.
2. For the indel score in the extension alignment, RepeatScout employed a linear gap penalty, whereas REPrise used an affine gap penalty (Gotoh, 1982), which is a more commonly used scoring system for sequence alignments.
3. In the masking step, RepeatScout masked all seeds within the identified repeats, whereas REPrise only masked the seed used for repeat detection. In other words, REPrise adopts a less stringent masking approach compared to RepeatScout, preserving more candidate regions for subsequent repeat detection.

In addition, we implemented REPrise to be applicable to long genome sequences. Our validation experiments showed that REPrise outperformed RepeatScout in the detection sensitivity of interspersed repeats using both rice and simulation genome datasets. We also applied REPrise to the complete human genome T2T-CHM13 and identified novel TE family candidates that have not yet been annotated.

## 2 Methods

### 2.1 Overview of the REPrise algorithm

Both REPrise and RepeatScout take genome sequences as inputs and output consensus sequences for each identified repeat families. The consensus sequence is a string composed of four characters A, T, C, G. Fig. 1 provides an overview of the REPrise and RepeatScout algorithms. Both algorithms first construct a seed table composed of seed sequences, their frequencies and locations in the input genome. As the seed sequences, RepeatScout uses *k*-mers that occur more than *c* times in the genome without allowing for substitutions. In contrast, REPrise allows for *d* substitutions. In accordance with a previous benchmark study (Ou *et al*., 2019), we set the value of *c* to 10. In the next step, the extension alignment is performed on the most frequent seeds in the seed table. Each alignment result from one seed sequence corresponds to one repeat family. While RepeatScout employs a linear gap scoring, REPrise uses an affine gap scoring. After the alignment, the genome regions corresponding to these seed sequences are masked, and the masked seeds are also removed from the seed table. REPrise adopts a less stringent masking approach to compared to RepeatScout. Then, the selection of most frequently occurring seed and the extension alignment is performed again. This cycle of the seed-and-extension and the masking is repeated until no more seeds are left in the seed table. Finally, REPrise further merges the consensus sequences of the identified repeat families using CD-HIT (Fu *et al*., 2012).

**Figure 1:**
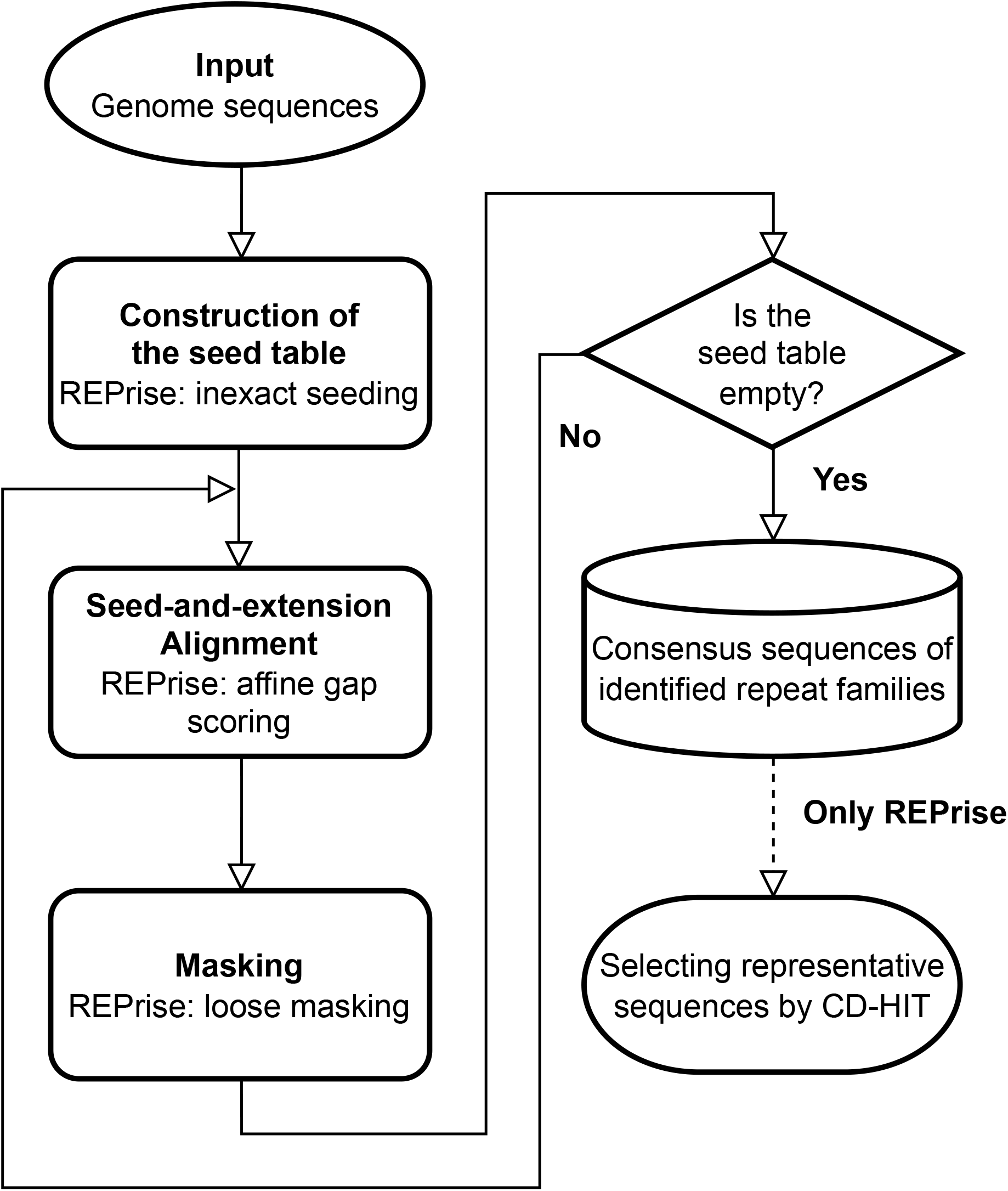
The schematic illustration of REPrise and RepeatScout algorithms. These algorithms first construct a seed table from the input genome sequences. REPrise utilizes inexact seeds, frequently appearing *k*-mers permitting *d* substitutions, for the table construction. Subsequently, these algorithms perform seed-and-extension alignments both ends of the seeds. REPrise adopts the affine gap socring in this step. These algorithms then mask seed regions in the detected repeat regions. REPrise performs looser seed masking than RepeatScout. This cycle of alignment and masking is repeated until the seed table is depleted. In REPrise, representative sequences are selected from the consensus sequences of repeat families using CD-HIT and the representative sequences are outputted.

### 2.2 Construction of the seed table

While RepeatScout constructs the seed table by scanning the genome only once, REPrise cannot adopt this approach because it employs inexact seeding. In addition, REPrise has to ensure that a *k*-mer is not counted multiple times from different seeds. To address these issues, REPrise use a suffix array to index the genome sequence *T*. The suffix array is a data structure that lists starting positions of all suffixes of a given string in the alphabetical order. It can be constructed with the time complexity of *O*(|*T* |) using the induced sorting algorithm (Nong *et al*., 2009) and is frequently used in bioinformatics sequence analysis software (Shrestha *et al*., 2014). By performing a binary search on the suffix array, for a given *k*-mer, we can count the frequencies of all *d*-similar *k*-mers in the genome. We defined two *k*-mers whose Hamming distance is less than or equal to *d* as *d*-similar *k*-mers.

REPrise constructs the seed table in the following manner:

1. The suffix array of the genome sequence is constructed.
2. For each *k*-mer seed, REPrise counts the frequencies of all *d*-similar *k*-mers in the genome. The count is taken as the frequency for each *k*-mer seed. In this step, Reprise allows that a k-mer is counted multiple times from different seeds. To accelerate this process, we employed parallel computations using multi-threading with OpenMP.
3. The *k*-mer seeds are sorted based on their frequencies.
4. In this sorted order, REPrise recounts the frequencies of all *d*-similar *k*-mers in the genome for each *k*-mer seed. In this recount step, REPrise does not count *k*-mer once counted again and thus can avoid multiple counts of the same *k*-mer from different *k*-mer seeds. Finally, only the seeds with a frequency of *c* or higher are retained in the seed table.

### 2.3 Seed-and-Extension alignment with affine gap scoring

The extension step of REPrise uses the banded alignment with affine gap scoring. The alignment procedure between two sequences is detailed in the Supplementary Material (Section S1). The consensus sequence *Q* among repeats is obtained by extending sequences from the seed region using the banded alignment. As the example, we introduce the rightward extension algorithm. Let *Q*_*t*_ denote the consensus sequence exteneded by *t* nucleotides to the right from the seed region. Given that the seed sequence is located at *M* positions in the genome, and the *M* sequences adjacent to the right of the seed region are defined as *S*_1_, *S*_2_, …, *S*_*M*_. The extension of the consensus sequence from *Q*_*t*_ to *Q*_*t*+1_ is performed according to the following equation:

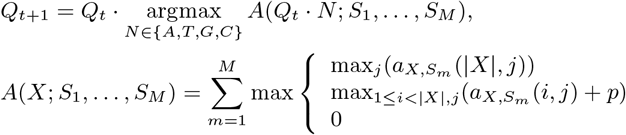

where · is an operator that appends a character to the right end of a sequence, *A*(*X*; *S*_1_, …, *S*_*M*_) is summation of the alignment score between *X* and each *S*_*m*_, and 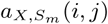 is the alignment score between between the first *i* characters of *X* and the first *j* characters of *S*_*m*_. The first term of *A* indicates that *X* aligns with *S*_*m*_ up to the end of *X*. The second term of *A* implys that *X* aligns only up to the middle of *S*_*m*_. *p* is an incomplete-fit penalty, which is set to -20, consistent with the value used in RepeatScout. The third term of *A* represents that *X* and *S*_*m*_ are not aligned at all.

The extension of *Q*_*t*_ is halted when *t*_*max*_ is not updated over a predefined length *w. t*_*max*_ is updated to *t* when the following conditions are satisfied:

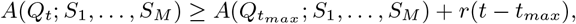

where *r* represents the penalty associated with the number of regions to be extended as the repeat regions in the genome and is set to three as with RepeatScout. When the extention is terminated, 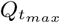 is concatenated to the right side of the seed sequence. In this study, *w* is set to 100 as with RepeatScout. The same extension process is performed to the left direction, and the final consensus sequence *Q* is obtained. As the left and right extensions are independent processes, they were computed in parallel by multi-threading with OpenMP.

### 2.4 Loose masking

Masking is a step to remove seed regions once used for repeat detection from the subsequent analysis, avoiding redundancy in repeat detection and reduceing the computation time. Specifically, the consensus sequence obtained in the extension step is re-aligned to the regions around the seed, and the seed sequences in the alignment regions are removed from the seed table (Supplementary Figure S2).

RepeatScout performs the realignment for all seed sequences within the consensus sequence. This approach effectively reduces the redundancy of repeat detection. However, because the realignment is conducted even for seeds not used for repeat detection, the detection sensitivity of different repeat families that share the same seed may be decreased. Therefore, REPrise adopts a loose masking approach, which targets only the seeds used for repeat detection. Unlike masking in RepeatScout, this masking approach prevents the loss of sensitivity from inappropriate masking but introduces the high redundancy among the detected repeat families. To mitigate this redundancy, after detecting all repeat families, REPrise performs CD-HIT on the identified repeat families and selects the representative sequences. According to the Wicker’s 80/80/80 rule for repeat detection (Wicker *et al*., 2007), we set the similarity threshold for CD-HIT at 80%.

### 2.5 Datasets and evaluation measures

We assessed the performance of REPrise using three genome datasets with curated repeat annotation: the rice, the simulation, and the complete human genome datasets. The first rice genome dataset (International Rice Genome Sequencing Project and Sasaki, 2005) has a manually curated repeat annotation using software tools such as RECON (Bao and Eddy, 2002) and serves as a standard benchmark for performance validation of repeat annotation software (Ou *et al*., 2019). The second simulation genome dataset was created using an existing TE insertion simulator that randomly incorporates sequences from a TE library into a long random sequence multiple times (Rodriguez and Makałowski, 2022). We created this dataset by adjusting the %_identity parameter of the simulator, which represents the sequence diversity across TE copies, ranging from 10 to 90. Specifically, for each inserted TE sequence, (50 – 0.5*%_identity)% of the sequence underwent mutation. We also adjusted the ratio of substitutions to indels for the mutation as follows: 100:0, 60:40 and 20:80. We used a default TE library comprising 20 TE families and inserted the TE sequences into a random sequence of 0.6 million lengths. This resulted in a sequence of approximately 4.6 million lengths for each parameter setting. The third complete human genome dataset is the T2T-CHM13 v2.0 (Nurk *et al*., 2022) genome with the repeat annotation (Hoyt *et al*., 2022). We used the thickStart and thickEnd columns in the .bed file for the repeat annotation. We also masked tandem repeats using TANTAN (Frith, 2011). Note that we did not mask tandem repeats in the rice genome dataset as the previous benchmark study did not use the masking tools.

We applied RepeatScout and REPrise to these genome datasets and identified the repeat families. Subsequently, we annotated the locations of the repeats within the genome using RepeatMasker and the identified repeat families. We then evaluated the performance of each software by comparing the software outputs with the curated annotation (Supplementary Figure S1(A)). We evaluated the overlap between the annotation and the software output at the nucleotide level and categorized the genomic regions into four groups: true positive (TP), true negative (TN), false positive (FP), and false negative (FN) (Supplementary Figure S1(B)). From these categories, we calculated four evaluation measures: sensitivity, specificity, precision, and F-score.

We also used the complete human genome dataset to discover novel interspersed repeat family candidates. We first mapped the detected repeat families by REPrise to the complete human genome using RepeatMasker. We then defined genome regions that met the following two criteria as novel repeat regions: (i) no overlap with tandem repeats, centromere satellites (Altemose *et al*., 2022), or segmental duplications (Vollger *et al*., 2022), and (ii) less than 20% overlap with genes (Pruitt *et al*., 2014) or known repeat regions. For each repeat family, if 40% or more of the mapped regions were classified as novel repeat regions, we labeled the repeat family as a novel repeat family.

## 3 Results

### 3.1 Evaluation in the rice genome dataset

We first explored the influence of seed length *k* on the repeat detection performance using the rice genome dataset. We set the gap open score as *o* = 5, the bandwidth as *b* = 5, and the gap extension score as *e* = 1 in this analysis. Supplementary Figure S3 shows the dependence of the detection performances on *k* of RepeatScout and REPrise with varying *d*. We found that the detection performance was highly dependent on *k* and the optimal *k* also depended on *d*. These findings underscore the importance of selecting appropriate *k* when using RepeatScout and REPrise. In this study, we selected the *k* with the highest sensitivity. Specifically, for the rice genome dataset analysis, we employed *k* = 15 for RepeatScout and REPrise with *d* = 0, *k* = 21 for REPrise with *d* = 1, and *k* = 25 for REPrise with *d* = 2. Furthermore, we assessed the effect of *b* and *e* on the detection performance (Supplementary Figure S4). We fixed *o* = 5 in this analysis. We found that *b* does not substantially affect the performance, and thus we set *b* = 5 because smaller *b* should speed up the computations. Moreover, we set *e* = 1 because this value was the best in all *e* when *b* = 5.

We next compared the repeat detection performance of RepeatScout and REPrise (Table 1). We found that REPrise exhibited the superior sensitivity and F-score compared to RepeatScout. In particular, REPrise outperformed RepeatScout even when employing the exact seed (*d* = 0). Since the improvement from the affine gap scoring was minor in this analysis (Supplementary Figure S4), the loose masking likely played a significant role in enhancing the performance. In addition, when evaluating the impact of *d* on the performance, the sensitivity increased with a rise in *d*, indicating that a larger *d* is advantageous for detecting novel repeats.

**Table 1:** Software performances for the rice genome dataset.

|  | Time[h:m:s] | Mem[GB] | Sensitivity | Specifiity | F-score |
| --- | --- | --- | --- | --- | --- |
| REPrise ($d = 0$) | 0:49:12 | 8.67 | 93.40 | 93.72 | **93.18** |
| REPrise ($d = 1$) | 1:42:55 | 14.49 | 93.51 | 93.46 | 93.10 |
| REPrise ($d = 2$) | 35:17:53 | 14.20 | **94.08** | 92.66 | 92.98 |
| RepeatScout | 0:52:17 | 5.49 | 89.62 | **95.82** | 92.23 |

The bold values are the highest scores among the software. "Mem" represents the amount of memory used. Note that RepeatScout is a single-threaded computation, whereas REPrise is a multi-threaded computation.

We also assessed the detection sensitivity across different repeat classes (Table 2). Our results showed that REPrise consistently outperformed RepeatScout for all repeat classes. In addition, the detection sensitivity of REPrise improved for all repeat classes as *d* increased. In particular, the performance largely improved for non-LTR elements, which were difficult to detect sensitively due to their high variability (Ou *et al*., 2019).

**Table 2:** Sensitivity results for each repeat class for the rice genome dataset.

|  | TIR | LTR | non-LTR | Helitron |
| --- | --- | --- | --- | --- |
| REPrise ($d = 0$) | 90.92 | 97.53 | 86.25 | 81.67 |
| REPrise ($d = 1$) | 91.16 | 97.55 | 87.22 | 81.20 |
| REPrise ($d = 2$) | **91.92** | **97.65** | **89.84** | **82.88** |
| RepeatScout | 85.20 | 96.39 | 76.32 | 73.05 |

The bold values are the highest scores among the software. We followed the repeat type classification of Ou *et al.* in this analysis (Ou *et al.* (2019)). TIR and LTR are the terminal inverted repeats and long terminal repeats, respectively.

We finally evaluated the program runtimes (Table 1). The computations were performed on an Intel Xeon Gold 6130 (16 cores) 2.1GHz CPU with 192 GB of memory. Note that REPrise was multi-threaded during both the seed table construction and extension steps, utilizing 16 and two threads, respectively. Even when using the same exact seeding method, REPrise with *d* = 0 is expected to take longer than RepeatScout because REPrise used the suffix array instead of the hash table to construct the seed table. However, their computational speeds were comparable, indicating that the parallelization by multi-threading in REPrise efficiently contributes to the speed-up. In addition, the runtimes increased as *d* increased, with *d* = 2 taking roughly 42 times longer than *d* = 0. We also analyzed how varying the number of threads influenced the runtime in REPrise (Supplementary Figure S5). We verified that increasing the numer of threads to 16 reduced the runtime, but there was no significant change in the runtime beyond 16 threads.

### 3.2 Evaluation in the simulation genome dataset

We next investigated the effects of the diversity among repeat sequences on the detection performance using the simulation datasets. We first compared the detection performance for various *k* values. Then, we selected *k* values as 13 for RepeatScout, and 13, 15, and 18 for REPrise with *d* values of 0, 1, and 2, respectively (Supplementary Figure S6).

Fig. 2 and Supplementary Figure S7 showed evaluation results for the simulation genome dataset. We verified that RepeatScout and REPrise had similar performances when the identity parameter was high, i.e. when the sequence similarity was high. Conversely, when the identity parameter dropped below 40, i.e. when the sequence similarity was low, REPrise with larger *d* showed higher performances. This result suggests that the inexact seeding was effective in detecting repeats that contain many mutations. Furthermore, when mutations were predominantly substitutions, there was no performance difference between REPrise (*d* = 0) and RepeatScout. However, when the majority of the mutations were indels, REPrise (*d* = 0) showed better performance than RepeatScout. This result indicates that the affine gap scoring and the loose masking improved the detection of repeats abundant in indels.

**Figure 2:**
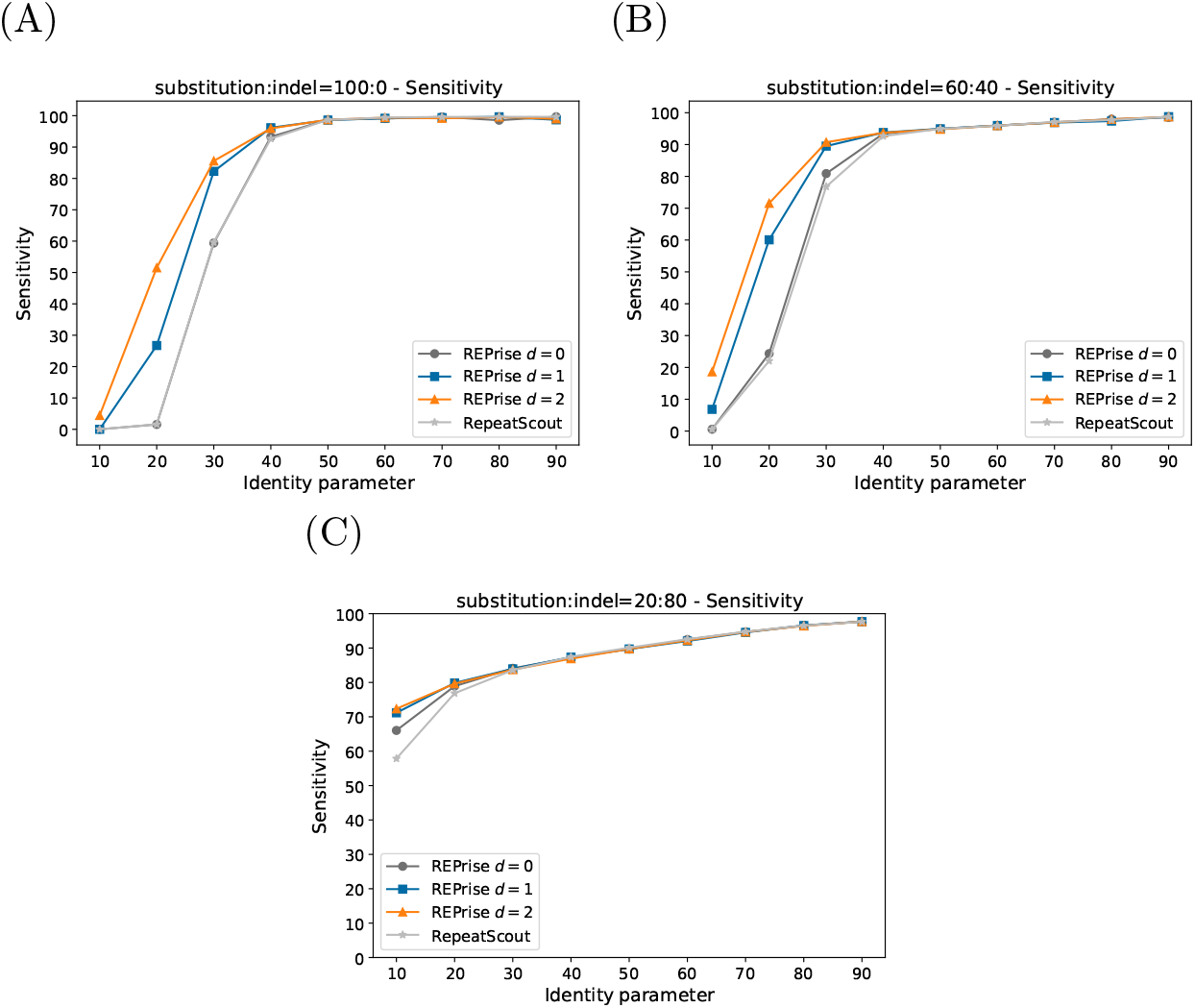
Dependence of the detection sensitivity of RepeatScout and REPrise on the identity parameter in the simulation genome dataset. The identity parameter represents the sequence diversity across TE copies. Specifically, for each inserted TE sequence, (50 – 0.5*identity parameter) of the sequence underwent mutation. The ratios of substitutions to indels were (A) 100:0, (B) 60:40, and (C) 20:80. The gray, black, blue, and orange lines represent RepeatScout, REPrise (*d* = 0, 1, 2), respectively. The x- and y - axes represent the identity parameter and the sensitivity, respectively.

We also found that the detection performance of these programs decreased with increasing substitution rates to indels when the identity parameter was low. This may be because the high substitution rate to indels prevented sharing of seed sequences among repeats in the same repeat family. Conversely, when the substitution rate to indels was lower, shared seed sequences among repeats in the same repeat family are likely to be retained. However, the low substitution rates (high indel rates) had a drawback that the number of repeat families was overestimated, because the indels fragment a single repeat family into multiple distinct repeat families (Supplementary Figure S7).

In all simulation datasets, the number of identified families increased as the identity parameter decreased, likely resulting from the fragmentation of detected repeat sequences. However, when the identity parameter was set extremely low, there was a noticeable decrease in the number of identified families, likely due to the reduced detection accuracy.

### 3.3 Analysis of the complete human genome dataset

We then applied REPrise to the complete human genome dataset. We selected 17, 21, and 23 as *k* for REPrise with *d* = 0, 1, and 2, respectively, after evaluating the detection performance among different *k* values (Supplementary Figure S8). Note that we could not employ RepeatScout for the human genome analysis because RepeatScout does not support application to long genomes. We validated that REPrise with *d* = 2 surpassed REPrise with *d* = 0 and *d* = 1 for all evaluation measures (Table 3).

**Table 3:** Software performances for the complete human genome dataset.

|  | Time[h:m:s] | Mem[GB] | Sensitivity | Specifiity | F-score |
| --- | --- | --- | --- | --- | --- |
| REPrise ($d = 0$) | 5:57:46 | 36.68 | 76.96 | 91.85 | 84.09 |
| REPrise ($d = 1$) | 37:40:56 | 67.64 | 77.68 | 92.11 | 84.63 |
| REPrise ($d = 2$) | 693:34:27 | 151.67 | **78.14** | **92.46** | **85.04** |

The bold values are the highest scores among $d$ . "Mem" represents the amount of memory used. The computations were performed on an Intel Xeon Gold 6148 Base 2.4GHz CPU with 32 parallel cores and 12GB memory per core.

We also explored the capability to identify novel repeat families for REPrise with *d* = 2. Consequently, we identified 17 novel repeat families (Supplementary Table S1). The novel repeat family with the highest proportion of novel repeat regions was novel_repeat_family-1 (NRF-1), where 100% of the repeat regions were novel. Upon investigation of these repeat regions, we found that they were located immediately upstream of each gene of the KRTAP9 gene family on chromosome 17 (Fig. 3). This result suggests that these repeat regions may be promoter regions that was duplicated during gene duplication in the KRTAP9 gene family. The novel repeat family with the second-highest proportion of novel repeat regions was NRF-2, with 71% of the 35 regions being novel. The majority of these regions were located on the Y chromosome, and parts of them were present in the long repetitive genomic regions newly identified by T2T-CHM13 (Supplementary Figure S9(A)). Given the abundance of tandem repeats on the Y chromosome, these regions may be part of tandem repeats, but they were not detected by existing tandem repeat detection tools, TRF (Benson, 1999) and WindowMasker (Morgulis *et al*., 2005) (Supplementary Figure S9(B)).

**Figure 3:**
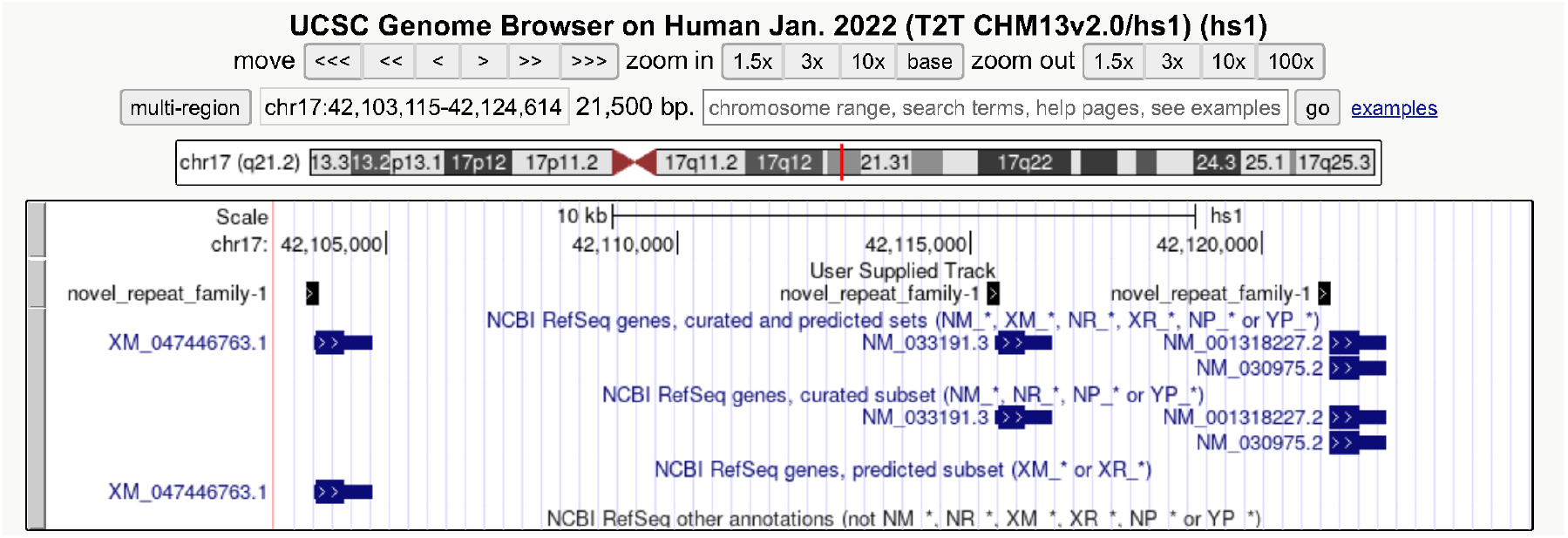
Genome regions of NRF1 (‘novel_repeat_family-1’ in the figure) with the NCBI RefSeq genes, displayed on the UCSC genome browser (Kent *et al*., 2002). NRF1 contains 9 repeat regions and 3 of the 9 regions are shown in this figure. Human readable names of XM_047446763.1 is KRTAP9-8, NM_033191.3 is KRTAP9-4, and NM_030975.2 and NM_001318227.2 are KRTAP9-9.

In the identified novel repeat families, two families may be unidentified subfamilies of existing LTR repeat families. The LTR regions are repeat regions that originally existed in pairs flanking the retroviral genome in the same orientation (Thomas *et al*., 2018). Although some LTR region are now isolated by the recombination, this characteristic LTR structure is retained in many genome regions. The first LTR subfamily candidate was NRF-10, which seemed to be a subfamily of the LTR19 family, because NRF-10 frequently appeared adjacent to fragments of the LTR19 family (Fig. 4(A,B)). In one case, NRF-10 was adjacent to the LTR-19 family while retaining the LTR-specific structure, suggesting that NRF-10 derived from the LTR (4(B)). To validate this hypothesis, we conducted a phylogenetic tree analysis of the LTR19 family and NRF-10. Specifically, we first extracted all annotated sequences of the LTR19 family and NRF-10 using SeqKit (Shen *et al*., 2016). When NRF-10 was found adjacent to the LTR19 family within a 500-base span, these sequences were concatenated into a single sequence using bedtools (Quinlan and Hall, 2010). We used the consensus sequence of MER50C as the outgroup of the phylogenetic tree ((Benachenhou *et al*., 2009)). These sequences were aligned with MAFFT version 7 (Katoh and Standley, 2013), and a rooted phylogenetic tree was constructed using trimAl (Capella-Gutiérrez *et al*., 2009) and IQ-TREE 2 (Minh *et al*., 2020). As a result, sequences containing NRF-10 form a subtree within the LTR19 family, suggesting that thse sequences including NRF-10 was a novel LTR repeat subfamily (Fig. 4(C)).

**Figure 4:**
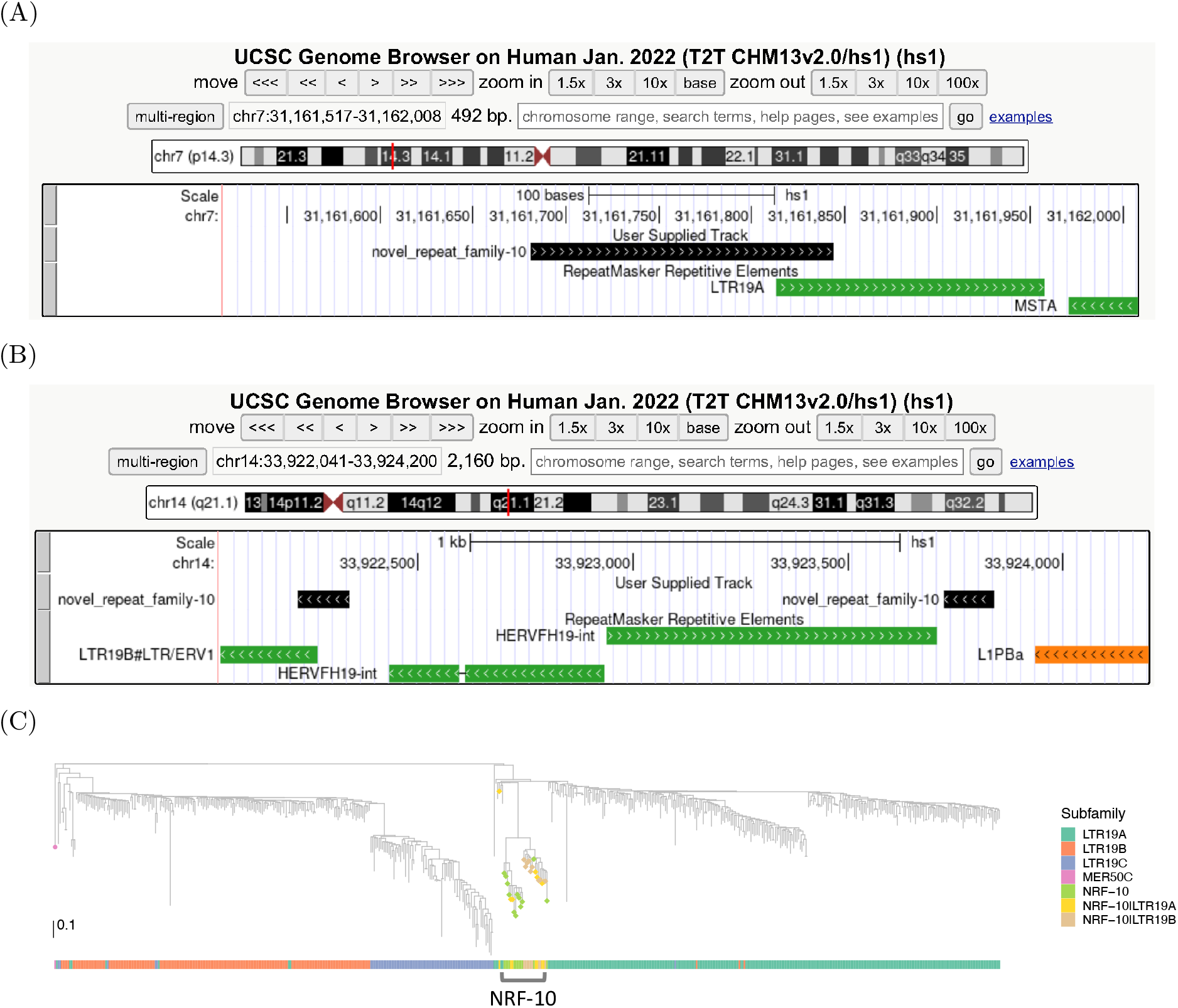
(A,B) Two examples of NRF-10 (‘novel_repeat_family-10’ in the figure) region with RepeatMasker Repetitive Elements displayed on the UCSC genome browser. (C) A phylogenetic tree of annotated regions of NRF-10 and LTR19 subfamilies. NRF-10|LTR19-A and NRF-10|LTR19-B are concatenated sequences of NRF-10 and LTR-19. The phylogenetic tree was visualized using ggtree (Xu *et al*., 2022) and ggtreeExtra(Xu *et al*., 2021).

The second LTR subfamily candidate was NRF-13, as many sequences from this repeat family overlapped with the LTR45 family (Supplementary Figure S10(A,B)). In addition, like NRF-10, there were also an example of LTR-specific structures forming in NRF-13 as well as NRF-10 (Supplementary Figure S10(C)). Therefore, we conducted a phylogenetic tree analysis for NRF-13. However, given the absence of a suitable outgroup for the LTR45 family, we established an unrooted phylogenetic tree(Supplementary Figure S11). We found that sequences encompassing NRF-13 also clustered into a distinct subtree, potentially representing another subfamily within the LTR45 family.

In this study, to investigate the usefulness of REPrise in identifying highly novel repeat families, we adopted strict criteria for their identification. Therefore, although we discovered some novel repeat families, we may have missed unidentified repeat subfamilies that are largely consistent with known repeat families but possess distinct regions. A more detailed analysis of individual repeat families detected by REPrise should reveal further novel repeat subfamilies.

## 4 Discussion

In this study, we developed REPrise, a highly sensitive tool for *de novo* detection of interspersed repeats. Although REPrise used a seed-and-extension approach similar to RepeatScout, REPrise enhanced sensitivity by incorporating three techniques: inexact seeding, affine gap scoring and loose masking. REPrise showed better repeat detection performance than RepeatScout on both empirical and simulation genome datasets, especially for repeats with many sequence mutations. Additionally, as a practical application of REPrise, we identified several novel repeat families in the complete human genome T2T-CHM13 v2.0.

The significant advantage of REPrise lies in its ability to apply sequence analysis across the entire length of gigantic genome sequences, such as the human genome. Currently, the widely used RepeatModeler2 pipeline for de novo repeat library construction relies on the internal execution of RepeatScout. RepeatModeler2 samples approximately 400MB of sequences during de novo repeat library creation, constituting only about 13% of the human genome. Consequently, there is a potential risk of overlooking repeat sequences in species with non-uniform compositions of interspersed repeats across regions and possessing exceptionally large genomes. In contrast, REPrise is inherently designed to handle large sequences and exhibits linear time complexity in relation to sequence length, addressing these limitations in current methodologies.

REPrise has multiple parameters, and the proper setting is crucial for achieving high detection performance. In particular, the seed length *k* has a significant impact on the detection performance. In this study, we applied REPrise with various *k* values to genome data and selected the *k* value that achieved the highest performance. However, in practical setting, *k* must be automatically determined. In RepeatScout, the default *k* was set using the formula: *k* = log_4_ |*T* |+1. This length was chosen such that the expected occurrence of a seed sequence within a random genome sequence is less than one. Based on this concept, we set the default *k* in REPrise as follows. Let *l* be the smallest natural number for which 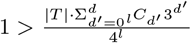, and then set *k* = *l* + 1. When comparing the automatically determined *k* value with the empirically determined *k* value, the two values were similar (Supplementary Table S2). This result suggests that our method of automatically determining *k* holds some level of validity. Exploring a more appropriate automatic method for determining *k* remains an essential topic for future research.

We envisioned two strategies for further enhancing the performance of REPrise. The first focuses on refining the seed design. Although the inexact seeding has improved detection sensitivity, the computational runtimes have also significantly increased. Therefore, improving computational speed is a critical issue. One possible solution is to adopt sparse seeding, a method commonly used in the sequence alignment, particularly for mapping long reads (Li, 2018). In this design, only a subset of subsequences in the genome serve as candidate seeds, rather than utilizing all subsequences. While this approach could drastically reduce the computation time, it may also diminish the detection sensitivity. Another direction for refining seed design is accounting for indels to further improve sensitivity. Recently, the ‘strobemer’ has been proposed as a seed design for efficient handling of indels in sequence alignment (Sahlin, 2021). The strobemer is a combination of multiple short *k*-mers and can account for potential insertions between two *k*-mers. The application of the strobemer to interspersed repeat detection may lead to the discovery of further novel repeat family candidates.

The second is enhancing the estimation accuracy of the number of repeat families. Our simulation experiments revealed that both RepeatScout and REPrise overestimated the number of families when the number of indels was high. Because many interspersed repeats in empirical genome have nested structures or large indels (Hoyt *et al*., 2022), the current estimates for the number of repeat families may be inflated. One feasible solution is to merge repeat families based on sequences surrounding the detected repeats. For instance, Red (Girgis, 2015) and RepLoc (Feng *et al*., 2020) determine repeat positions in the genome using *k*-mers without identifying the repeat families, and integrating these tools into REPrise could improve the accuracy of family number estimates. Another approach is the use of graph genome methods, which have gained attention for representing genetic variation among individuals (Eizenga *et al*., 2020). The sequence alignment tools in this paradigm have already been developed (Garrison *et al*., 2018), and their application to interspersed repeat detection is an emerging area (Groza *et al*., 2023).

## Acknowledgements

We thanks to members of Hamada Laboratory, especially Kento Kubo and Masato Kosuge for their valuable comments. A.T. is grateful to Sigehiro Kuraku (National Institute of Genetics) for insightful discussions. A.T. thanks to Naoki Konno (Department of Biological Science, Graduate School of Science, The University Tokyo) for technical supports in phylogenetic tree creation. Computations were partially performed on the NIG supercomputer at ROIS National Institute of Genetics.

