## Supplemental File for "REPrise: *de novo* interspersed repeat detection using inexact seeding"

### Supplementary Materials for "REPrise: *de novo* interspersed repeat detection using inexact seeding"

### S1 Supplementary Text

In the alignment of REPrise extension step, for two sequences $X$ and $Y$, three dynamic programming (DP) tables $M$, $I_X$ and $I_Y$ are filled according to the following update formula:

$$M(i,j) = \max \begin{cases} M(i-1,j-1) + \mu_{ij} \\ I_X(i-1,j-1) + \mu_{ij} \\ I_Y(i-1,j-1) + \mu_{ij} \end{cases}$$

$$I_X(i,j) = \max \begin{cases} M(i-1,j) + o \\ I_X(i-1,j) + e \end{cases}$$

$$I_Y(i,j) = \max \begin{cases} M(i,j-1) + o \\ I_Y(i,j-1) + e \end{cases}$$

, where $1 \leq i \leq |X|$, $1 \leq j \leq |Y|$ and $i-b \leq j \leq i+b$. Here, $\mu_{ij}$ denotes the score based on either a match or a mismatch between $i$-th charater of $X$ and $j$-th character of $Y$. The score takes $+1$ (for the match) or $-1$ (for the mismatch). $o$ is a gap open score, $e$ is a gap extension score, and $b$ is a banded width parameter. We set the initial values of the DP table as follows: $M(0,0) = 0$, $I_X(0,0) = I_Y(i,j) = -\infty$, $M(i,0) = M(0,j) = -\infty$ and $I_X(i,0) = I_Y(0,i) = o + i * e$ $(1 \leq i \leq b)$. The alignment score $a(i,j)$ is obtained as $\max(M(i,j), I_X(i,j), I_Y(i,j))$.

In the masking step, REPrise performs pairwise alignment between the consensus sequence $Q$ and the sequences surrounding the region matching the seeds. Specifically, the extension alignment is performed from the endpoint of the seed. As the example, we introduce the rightward extension algorithm. The sequences $S_1, S_2, \ldots, S_M$ are defined as the $M$ sequences adjacent to the right of the seed region. The alignment score $a_{Q,S_m}(i,j)$ represents the alignment score between the first $i$ characters of $Q$ and the first $j$ characters of $S_m$. Let $(i', j')$ denote the pair of $(i,j)$ that satisfy $\max a_{Q,S_m}(i,j)$. REPrise masks the region corresponding to the first $j'$ characters on $S_m$. Similarly, REPrise performs masking in the left direction. Additionally, REPrise masks the matched seed of the starting point of $S_m$.

#### S2 Supplementary Figure

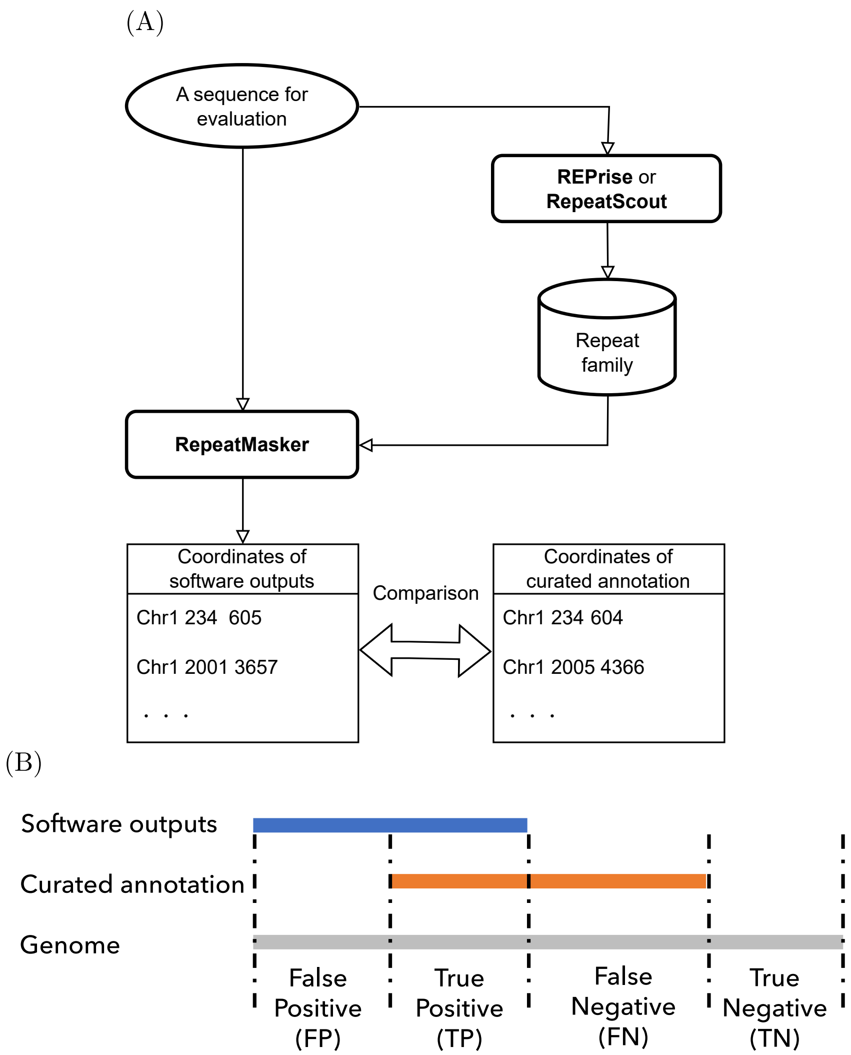


Figure S1: The schematic illustration of the evaluation method of REPrise and RepeatScout. **(A)** The flowchart of the evaluation. The output repeat families by REPrise and RepeatScout were located in the input genome using RepeatMasker, and the repeat positions were then compared to the repeat positions of the curated annotation. **(B)** The genome regions were divided into four parts, TP, TN, FP and FN, based on the overlap between the software output and the curated annotation. The evaluation measures were calculated from the number of bases in the four regions.

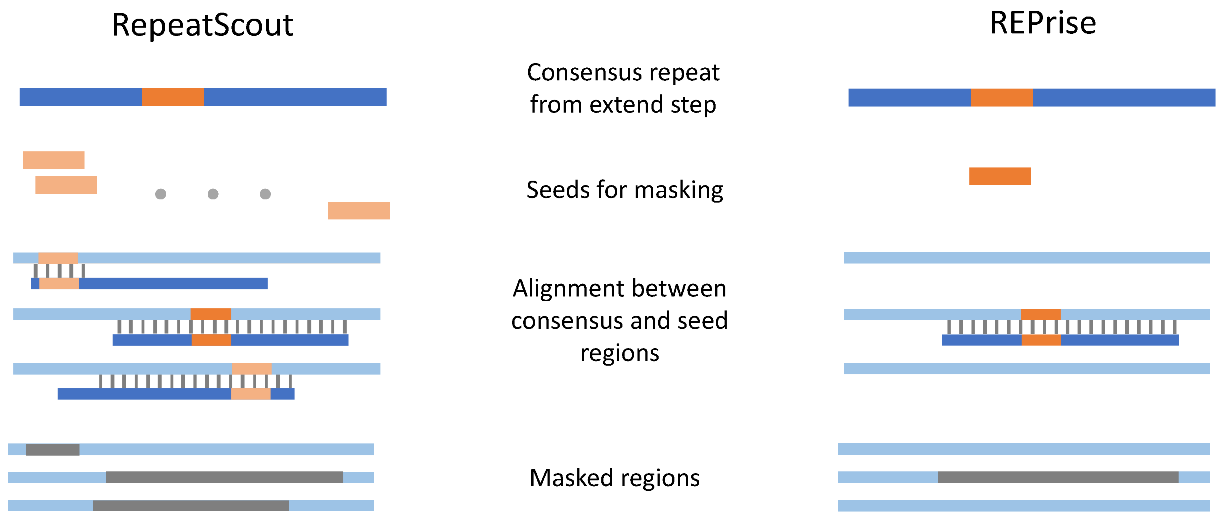


Figure S2: The schematic illustration of masking differences between RepeatScout and REPrise. The blue line represents the consensus repeat determined by the extension step, the dark orange box represents the seed that was the starting point of the extension step and the light orange box represents the all seeds in the consensus repeat. REPRise only uses the seeds in the dark orange box for masking, whereas RepeatScout uses the seeds in both the dark and light orange boxes for masking. In addition, the gray vertical bars represents aligned regions, the light blue line represents the genome sequence, and the gray line represents the finally masked regions.

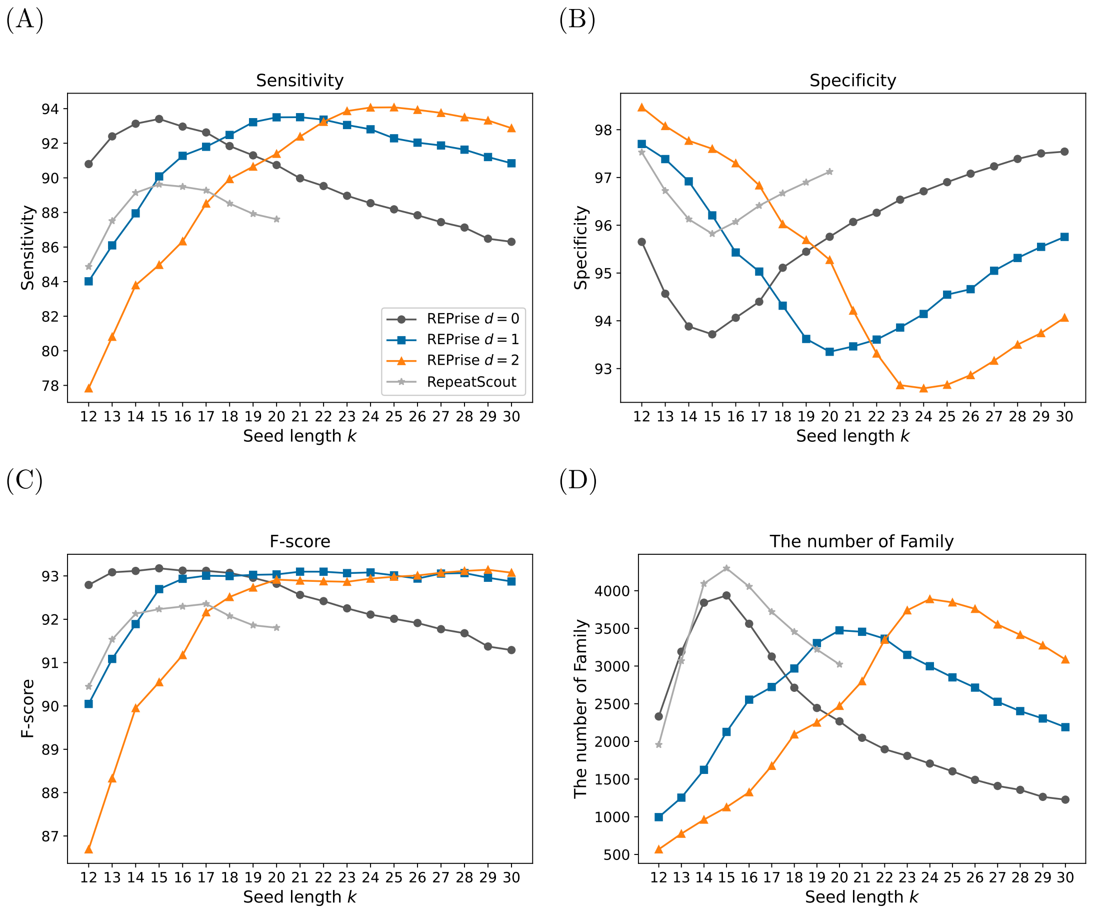


Figure S3: Dependence of the evaluation measures of RepeatScout and REPrise on the seed length $k$ in the rice genome dataset. The gray, black, blue, and orange lines represent RepeatScout, REPrise ($d = 0, 1, 2$), respectively. The x-axis represents the seed length $k$. The y-axis represents (A) the sensitivity, (B) the specificity, (C) the F-score and (D) the number of families.

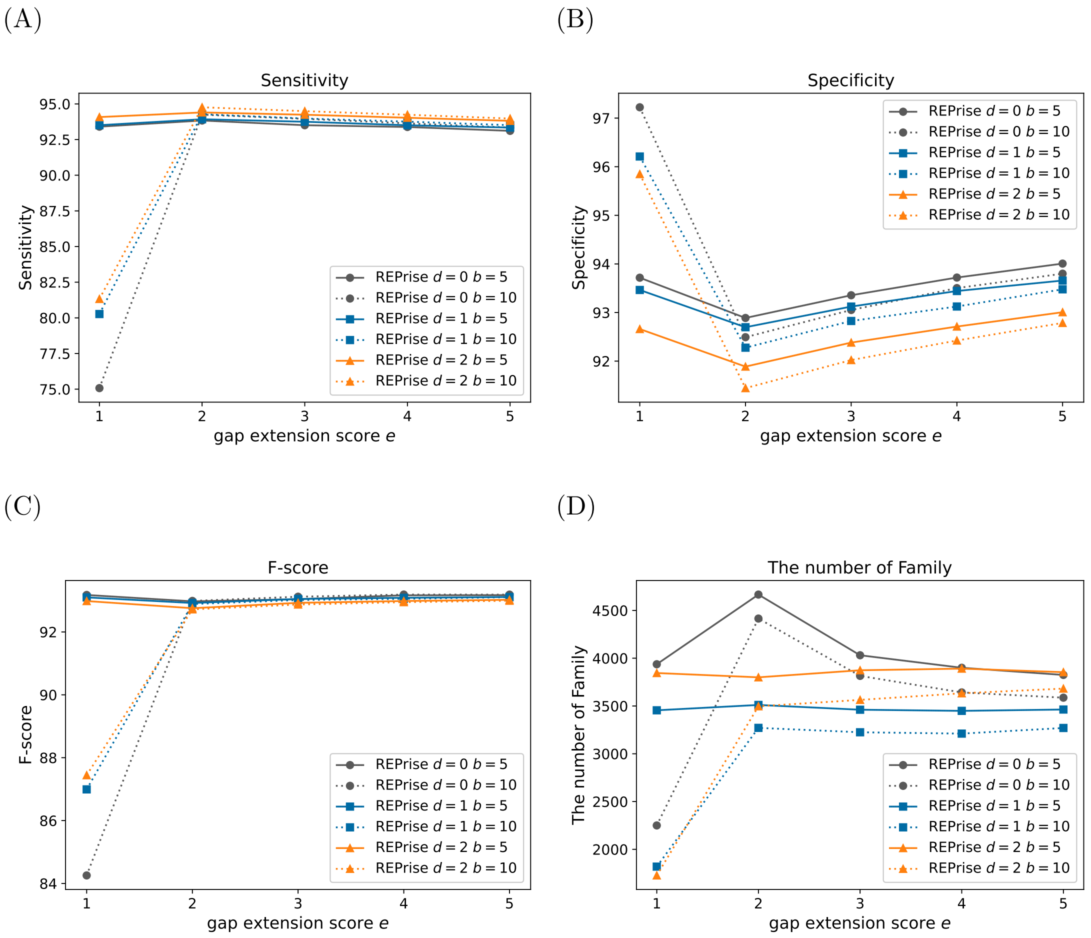


Figure S4: Dependence of the evaluation measures of REPrise on the banded witdh $b$ and the gap extension score $e$ in the rice genome dataset. The black, blue and orange lines represent REPrise with $d = 0, 1, 2$, respectively. The solid and dashed lines represent REPrise with $b = 5, 10$, respectively. The x-axis represents the gap extension score $e$. The y-axis represents (A) the sensitivity, (B) the specificity, (C) the F-score and (D) the number of families.

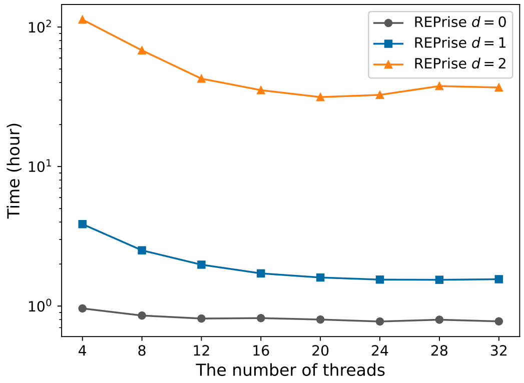


Figure S5: Dependence of the runtimes of REPrise on the number of threads for pallalel computation. The black, blue and orange lines represent REPrise with $d = 0, 1, 2$, respectively. The x- and the y- axes represents the number of threads and the runtimes, respectively. The time is shown in logarithmic scale.

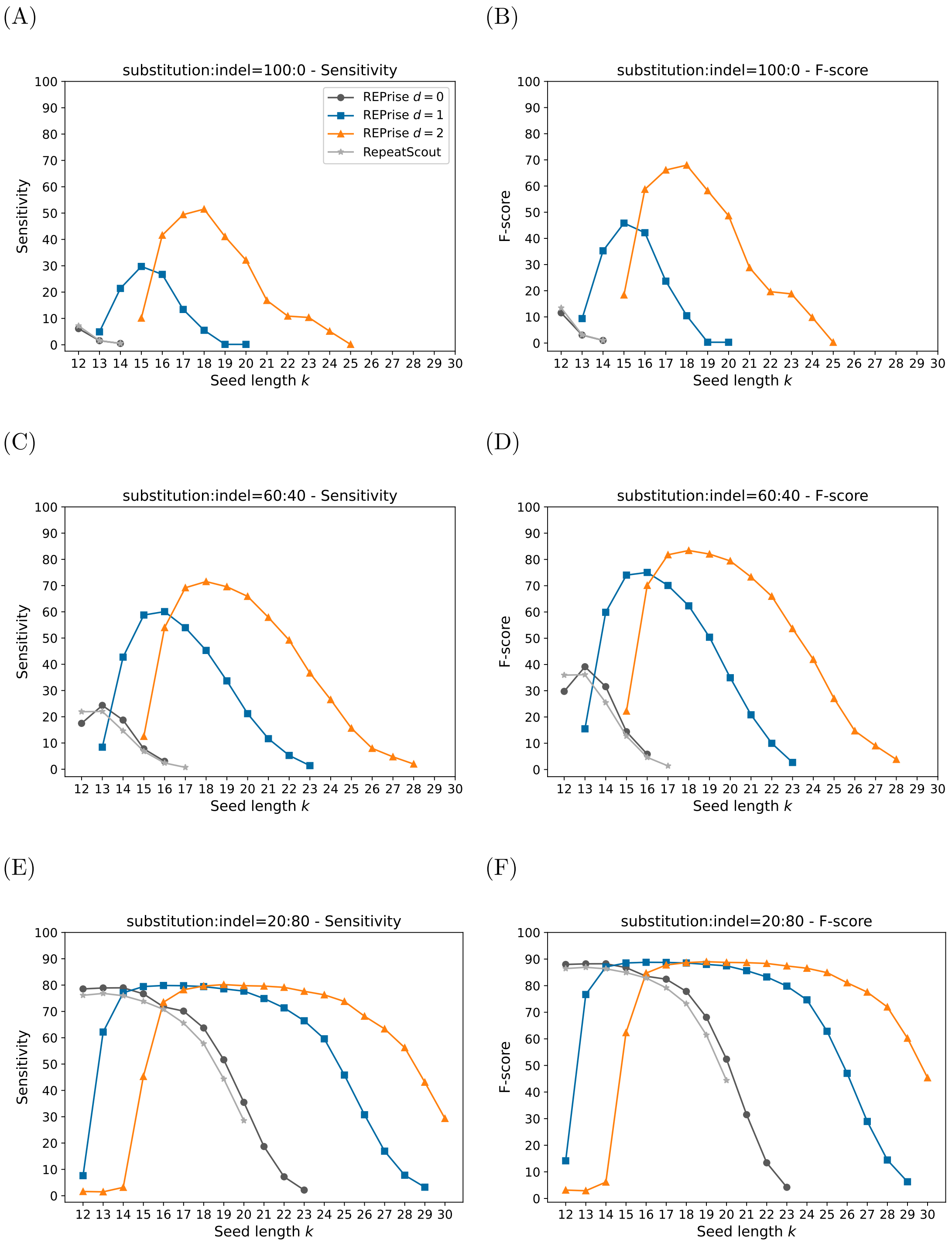


Figure S6: Dependence of the evaluation measures of RepeatScout and REPrise on the seed length $k$ in the simulation genome dataset. The gray, black, blue, and orange lines represent RepeatScout, REPrise ($d = 0, 1, 2$), respectively. The x-axis represents the seed length $k$. The y-axis represents (A)(C)(E) the sensitivity and (B)(D)(F) the F-score. We set the identity parameter to 20 and the ratio of substitutions to indels to (A)(B) 100:0, (C)(D) 60:40 and (E)(F) 20:80.

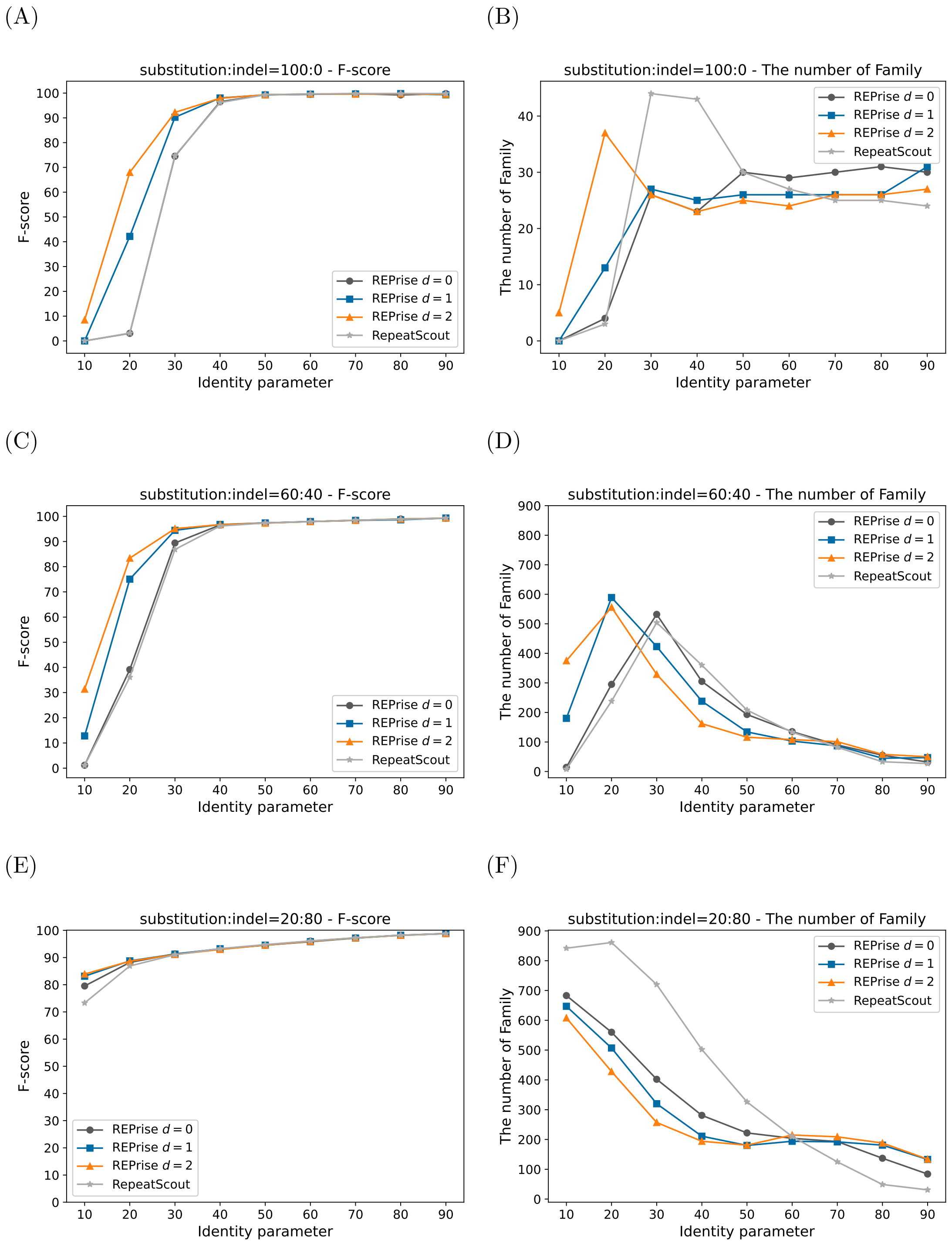


Figure S7: Dependence of the F-score and the number of families of RepeatScout and REPrise on the identity parameter in the simulation genome dataset. The ratios of substitutions to indels were (A)(B) 100:0, (C)(D) 60:40, and (E)(F) 20:80. The gray, black, blue, and orange lines represent RepeatScout, REPrise ($d = 0, 1, 2$), respectively. The x-axis represents the identity parameter. The y-axis represents (A)(C)(E) the F-score and (B)(D)(F) the number of families.

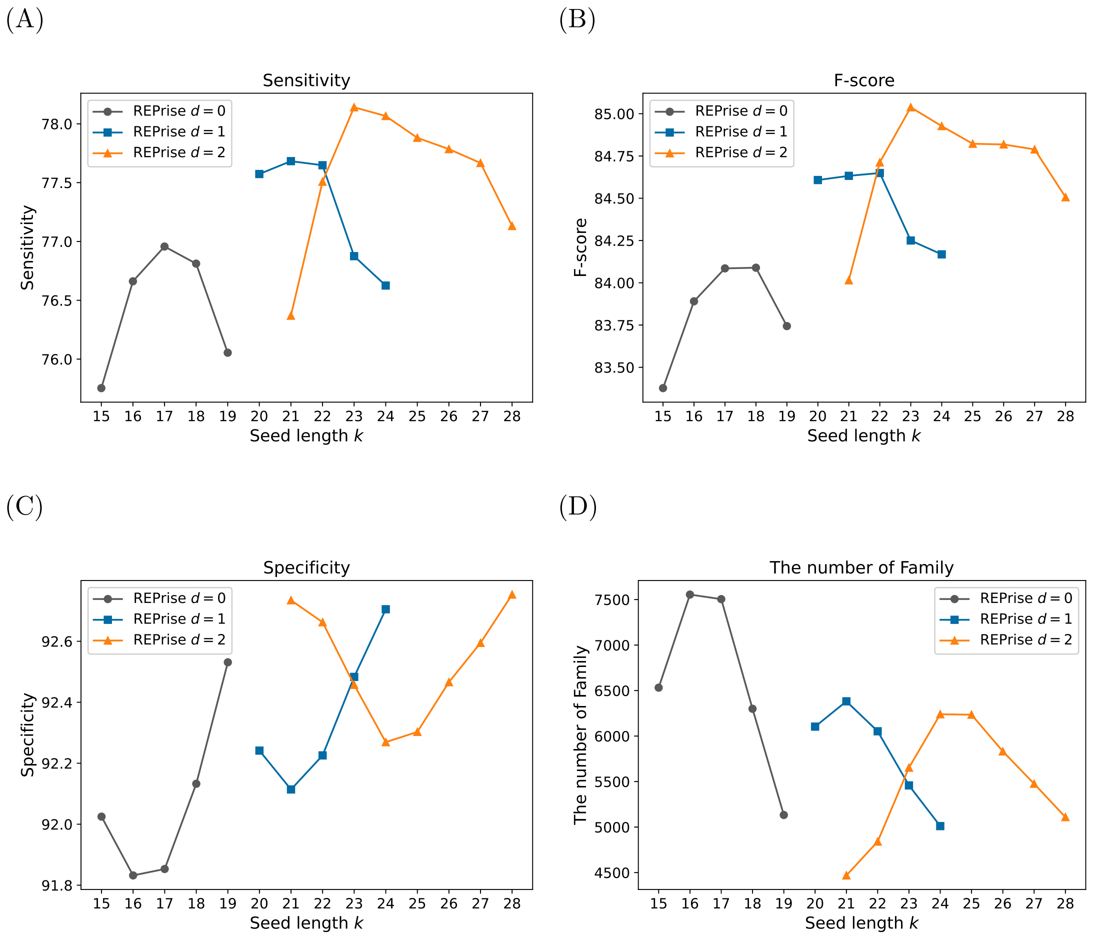


Figure S8: Dependence of the evaluation measures of RepeatScout and REPrise on the seed length $k$ in the complete human genome dataset. The black, blue, and orange lines represent REPrise with $d = 0, 1, 2$, respectively. The x-axis represents the seed length $k$. The y-axis represents (A) the sensitivity, (B) the specificity, (C) the F-score and (D) the number of families.

(A)

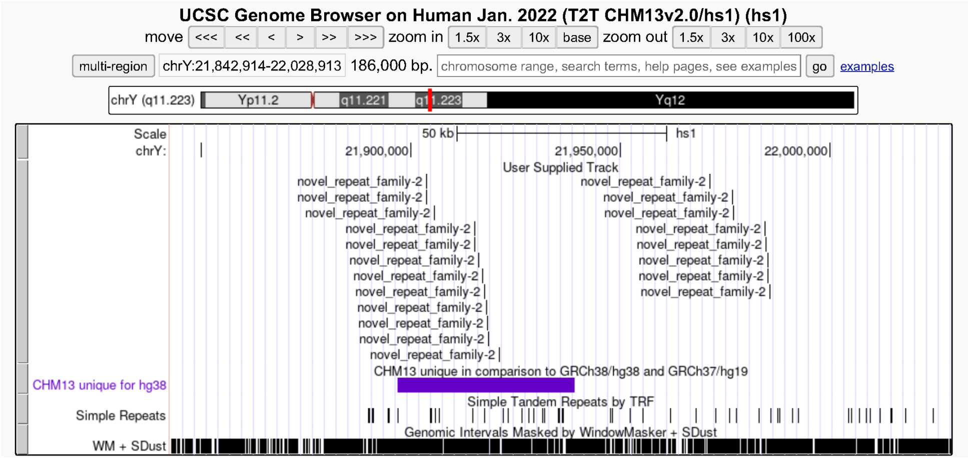


(B)

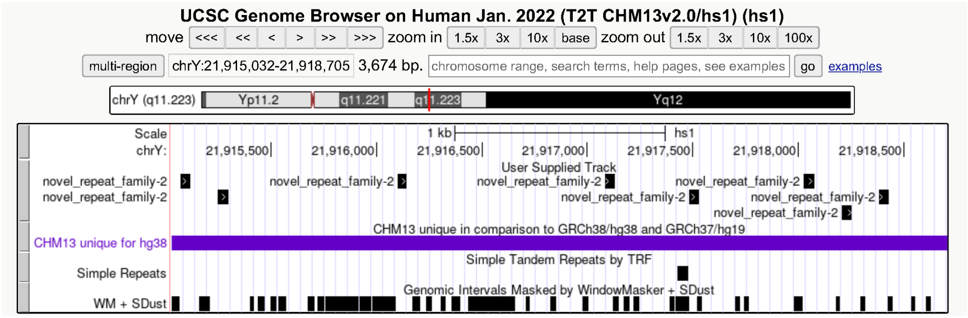


Figure S9: (A)(B)Genome regions of NRF2 from the UCSC genome browser.

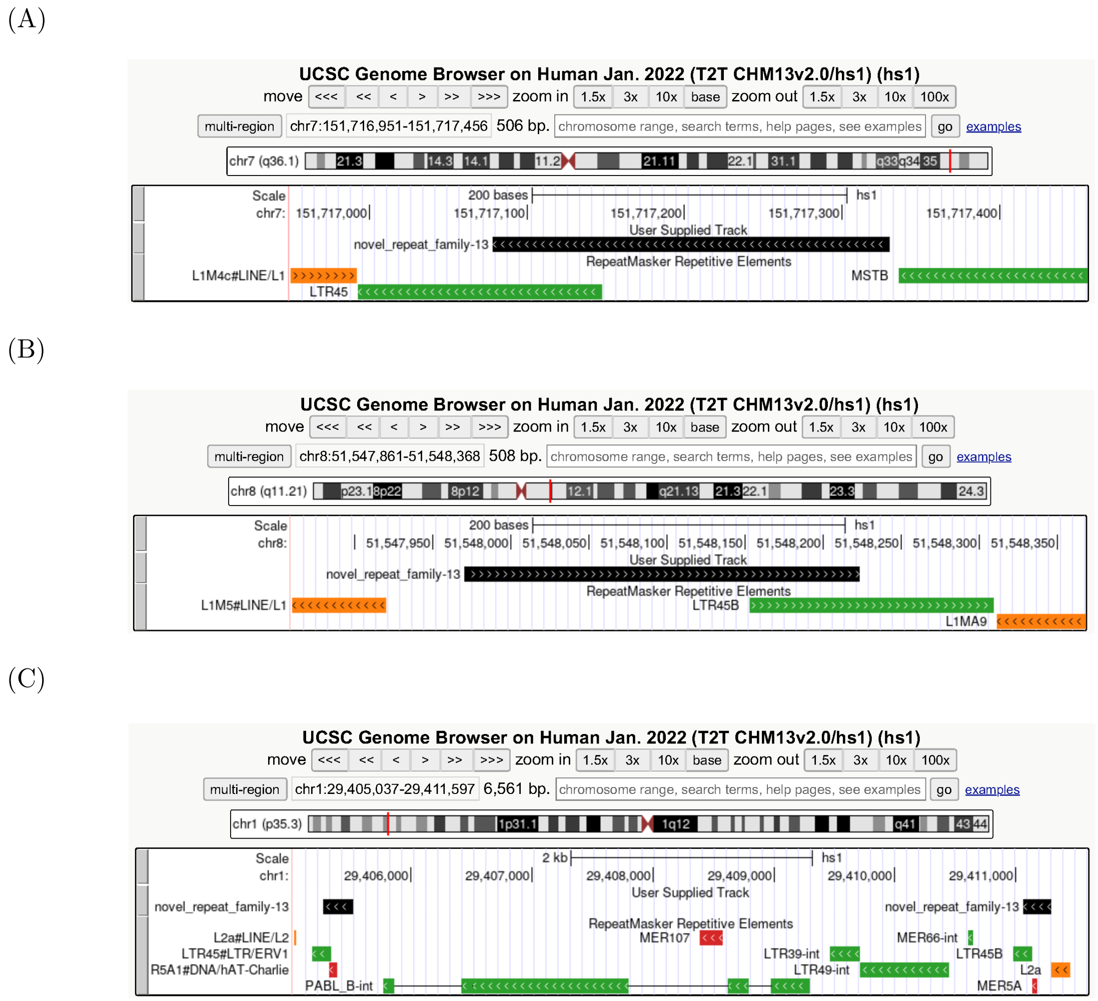


Figure S10: Genome regions of NRF13 from the UCSC genome browser.

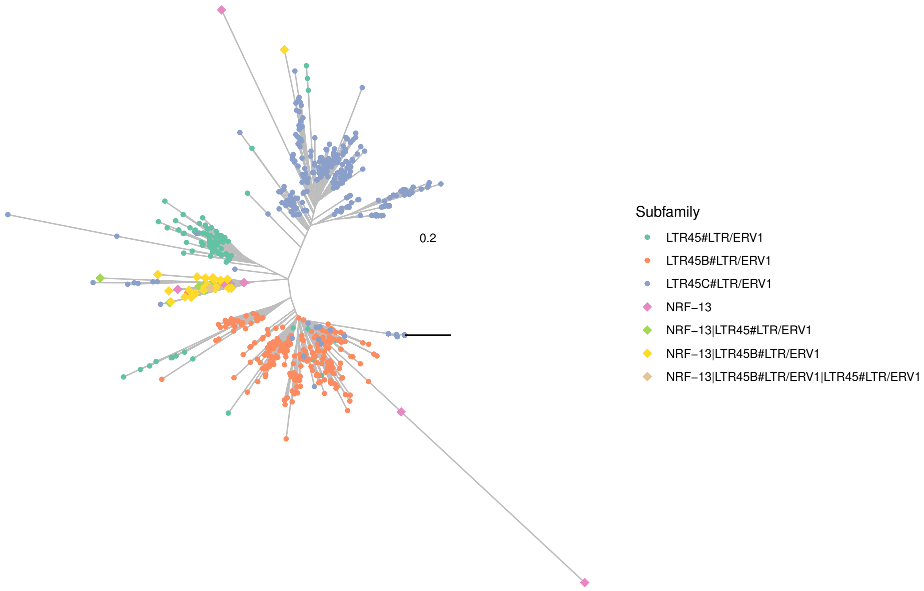


Figure S11: A phylogenetic tree of annotated regions of NRF-13 and LTR45 subfamilies. "|" indicates merge of two sequences.

### S3 Supplementary Table

Table S1: Novel repeat families detected by REPrise with $d = 2$

| Family name | Consensus sequence length | The number of NRRs | Total number of repeat regions | Percentages of the NRRs[%] |
|---|---|---|---|---|
| NRF-1 | 211 | 9 | 9 | 100.00 |
| NRF-2 | 50 | 25 | 35 | 71.43 |
| NRF-3 | 215 | 33 | 57 | 57.89 |
| NRF-4 | 145 | 9 | 16 | 56.25 |
| NRF-5 | 824 | 32 | 57 | 56.14 |
| NRF-6 | 365 | 25 | 46 | 54.35 |
| NRF-7 | 92 | 7 | 13 | 53.85 |
| NRF-8 | 486 | 17 | 32 | 53.12 |
| NRF-9 | 107 | 18 | 34 | 52.94 |
| NRF-10 | 164 | 14 | 31 | 45.16 |
| NRF-11 | 794 | 9 | 20 | 45.00 |
| NRF-12 | 60 | 29 | 65 | 44.62 |
| NRF-13 | 250 | 18 | 41 | 43.90 |
| NRF-14 | 120 | 36 | 84 | 42.86 |
| NRF-15 | 433 | 11 | 26 | 42.31 |
| NRF-16 | 581 | 19 | 46 | 41.30 |
| NRF-17 | 73 | 75 | 187 | 40.11 |

The NRFs and NRRs are abbreviations of the novel repeat families and novel repeat regions, respectively. We defined NRRs as genome regions that met the following two criteria (i) no overlap with tandem repeats, centromere satellites , or segmental duplications, and (ii) less than 20% overlap with genes or known repeat regions. Total number of repeat regions is counts of all repeat regions in the repeat family, and the NRF is the repeat family satisfying
(Number of NRRs/Total number of repeat regions) $\geq 0.4$.

Table S2: Comparison of empirically and automatically determined seed lengths.

| | Rice $d = 0$ | Rice $d = 1$ | Rice $d = 2$ | Sim $d = 0$ | Sim $d = 1$ | Sim $d = 2$ | T2T $d = 0$ | T2T $d = 1$ | T2T $d = 2$ |
|---|---|---|---|---|---|---|---|---|---|
| Empirical | 15 | 21 | 25 | 13 | 16 | 18 | 17 | 21 | 23 |
| Automatic | 16 | 19 | 21 | 14 | 17 | 19 | 17 | 20 | 23 |

"Sim" represents the seed length for the simulation dataset whose identity parameter was 60 and substitution:indel was 60:40. Because the seed length vary depending on the parameters of the simulation dataset, we presented the typical seed lengths in this table.
